# Resident Synovial Macrophages in Synovial Fluid: Implications for Immunoregulation in Infectious and Inflammatory Arthritis

**DOI:** 10.1101/2023.09.29.560183

**Authors:** Karen I. Cyndari, Breanna M. Scorza, Zeb R. Zacharias, Leela Strand, Kurayi Mahachi, Juan Marcos Oviedo, Lisa Gibbs, Danielle Pessoa-Pereira, Graham Ausdal, Dylan Hendricks, Rika Yahashiri, Jacob M. Elkins, Trevor Gulbrandsen, Andrew R. Peterson, Michael C. Willey, Keke C. Fairfax, Christine A. Petersen

## Abstract

**Objectives:** Resident synovial macrophages (RSM) provide immune sequestration of the joint space and are likely involved in initiation and perpetuation of the joint-specific immune response. We sought to identify RSM in synovial fluid (SF) and demonstrate migratory ability, in additional to functional changes that may perpetuate a chronic inflammatory response within joint spaces.

**Methods:** We recruited human patients presenting with undifferentiated arthritis in multiple clinical settings. We used flow cytometry to identify mononuclear cells in peripheral blood and SF. We used a novel transwell migration assay with human *ex-vivo* synovium obtained intra-operatively to validate flow cytometry findings. We used single cell RNA-sequencing (scRNA-seq) to further identify macrophage/monocyte subsets. ELISA was used to evaluate the bone-resorption potential of SF.

**Results:** We were able to identify a rare population of CD14^dim^, OPG^+^, ZO-1^+^ cells consistent with RSM in SF via flow cytometry. These cells were relatively enriched in the SF during infectious processes, but absolutely decreased compared to healthy controls. Similar putative RSM were identified using *ex vivo* migration assays when MCP-1 and LPS were used as migratory stimulus. scRNA-seq revealed a population consistent with RSM transcriptionally related to CD56^+^ cytotoxic dendritic cells and IDO^+^ M2 macrophages.

**Conclusion:** We identified a rare cell population consistent with RSM, indicating these cells are likely migratory and able to initiate or coordinate both acute (septic) or chronic (autoimmune or inflammatory) arthritis. RSM analysis via scRNA-seq indicated these cells are M2 skewed, capable of antigen presentation, and have consistent functions in both septic and inflammatory arthritis.

## Introduction

Damage to the articular surface of joints resulting in arthritis may be secondary to infection, inflammation, and chronic or acute trauma. Different etiologies of arthritis result in unique local immune environments within the joint space. While healthy synovium delineates an immune-privileged space to which few circulating cells gain entry (1), there are significant numbers of immune cells in the synovial fluid (SF) of pathologic joints (2; 3; 4). This indicates the localized synovial immune response is coordinated by the synovium itself, which limits entry to the SF by actively sequestering inflammatory damage (5) versus allowing circulating immune cells to enter SF.

Synovium is primarily composed of two types of cells: Fibroblast-like Synoviocytes (FLS) and Resident Synovial Macrophages (RSMs), also known as Type A cells. FLS express MHC Class II and produce lubricating joint fluid, including hyaluronan (6). In pathologic settings, FLS are involved in joint inflammation and, ultimately, cartilage destruction (6). Conversely, RSMs compose approximately 10% of the synovium and have only been identified using tissue histology to date. RSMs are described as constitutively anti-inflammatory, as opposed to circulating monocytes which may either assist with sequestration of pathology or provide further momentum toward significant cellular collateral damage (5).

It was recently discovered that RSMs are derived from embryonic precursor cells and perpetuate through self-proliferation within synovium (7). Certain cellular surface receptors, chemokines, or structural proteins such as CD68, osteoprotegerin (OPG/TNFRSF11B), CX3CR1, ZO-1/TJP1, F11R/JAM-A/JAM-1/CD321, and Triggering Receptor Expressed on Myeloid cells 2 (TREM2) (8; 7) have all been proposed to identify these cells, yet it remains difficult to differentiate RSMs from macrophages recruited from circulation. RSMs have not been further evaluated in SF as it is unclear if they can migrate out of tissue. As inflammation dysregulates the tight junctions connecting these epithelial-like RSMs (7), it is possible that these cells could leave the synovium and participate in joint space immune responses.

Here, we used flow cytometry to describe a subset of CD14^dim^OPG+ZO-1+ M2 macrophages enriched in SF of pathologic joints consistent with previously published histological descriptions of RSMs. *Ex vivo* migration experiments validated migration from tissue. Single cell RNA sequencing (scRNA-seq) reveals these putative RSMs had dysregulated complement in settings of inflammatory arthritis, and a unique reactome signature involving threonine, niacin, and thiamine metabolism. This work is important in understanding how damage to the joint space is initiated and perpetuated both during infectious and inflammatory arthritis.

## Materials and Methods

### Patient Recruitment

These studies were approved by the Institutional Review Board (IRB) at the University of Iowa Hospitals and Clinics. For SF studies, we recruited patients under evaluation for septic arthritis. Exclusion criteria included significant joint trauma, joint surgery, or immunomodulatory medications. All patients provided a blood sample. Not all patients required or had successful arthrocentesis. SF samples were classified as Normal, Non-Inflammatory, Inflammatory, or Septic based on established guidelines (9). For synovium procurement, 6 patients >18 years of age receiving a scheduled, non-emergent, total joint replacement or resection arthroplasty secondary to infection were recruited. Synovium (knee) or pulvinar (hip) was sterilely obtained during normal operating protocol.

### PBMC and SF Cell Sample Preparation

Peripheral blood mononuclear cells (PBMCs) were isolated from whole blood over Ficoll-Paque PLUS (Fisherbrand). For SF samples, a 200 µl aliquot was centrifuged at 1000 RCF for 10 minutes and stored at -80°C for ELISA. The remainder of the SF was treated with bovine testes hyaluronidase (Sigma-Adritch) according to manufacturer’s SF clarification protocol. SF was then filtered through a 70µm nylon mesh strainer (Fisher Scientific), diluted to 10 mL with PBS, and centrifuged at 400 RCF for 10 minutes at room temperature. Supernatant was discarded. SF cells (SFCs) and PBMCs were counted and then cryopreserved in 90% FBS with 10% DMSO.

### Transwell Migration Assay

Synovium (knee) or pulvinar (hip) acquired in the operating room was transferred to the laboratory in PBS on ice. Whole synovium was washed twice in PBS, then sterilely dissected into 4×4 mm segments, and washed again. Segments were placed into 24-well transwell inserts with 5 µm pores (Corning), a size that should allow monocyte and macrophage migration, but prevent fibroblast migration (10). Bottom wells were treated with LPS (1, 10, or 100 µg/mL, Sigma-Aldrich) or MCP-1 (25 ng or 250 ng/mL, Fischer Scientific) in DMEM supplemented with 10% FBS and 1% penicillin/streptomycin. The inserts were then placed in the well, and enough media to cover the tissue was placed in the insert (approximately 200-400 µl). The transwell plates were incubated for 24 hours at 37°C and 5% CO_2_. At day 1, 2, 3, 5, and 7, changes to the synovium were compared to *in vivo* controls also obtained intra-operatively (**Suppl. Figure 2**). H&E staining was performed to evaluate changes to the synovial intimal and sub-intimal lining with MCP-1 and LPS treatment (**Suppl. Figure 3**) Validation for tissue survival was performed out to 7 days with normoxic (21% oxygen) and hyperoxic (50% oxygen) conditions and Caspace-3 immunohistochemistry (**Suppl. Figure 4**). After incubation, the cells in the bottom well media were counted, and a crystal violet assay was performed on cells adhered to the bottom of the well and the underside of the transwell per standard protocol. Migratory cells in solution were analyzed using flow cytometry.

### Flow Cytometry Analysis

Samples were plated in a 96-well round-bottom plate for single stain, unstained, patient test samples, and/or Fluorescence Minus One (FMO) controls. Controls were plated at 2×10^5^ cells per well, and patient test samples at 1-2×10^6^ cells per well. Antibodies are listed in **Suppl. Table 1**. For intracellular staining (OPG, CD68, F11R, TREM2, ZO-1 and RANKL), cells were fixed and permeabilized with Fixation Buffer and Intracellular Staining Perm Wash Buffer (BioLegend) or Cytofix/Cytoperm Fixation/Permeabilization Solution Kit (BD Bioscences). Final flow cytometry gating strategy for SFCs and PBMCs shown in **Suppl. Figure 1**. Flow cytometry was performed on a Cytek Aurora cytometer (Bethesda, MD). Analysis was performed using FlowJo (Ashland, OR) software.

### Enzyme-linked Immunosorbent Assays (ELISA)

Human TIMP-1 (RAB0466-1KT), TGF-BETA (RAB0460-1KT), TRACP (RAB1755-1KT), OPG/TNFRSF11B (RAB0484-1KT), IFN GAMMA (RAB0222-1KT), TNF-ALPHA (RAB0476-1KT), MMP-9 (RAB0372-1KT) and BMP2 (RAB0028-1KT) ELISA kits were obtained from Millipore Sigma, and sRANKL (MBS262624) kit from MyBioSource. For TIMP-1 and MMP-9, SF was diluted 1:500, for sRANKL dilution was 1:100, and all others were diluted 1:20. TGF-β1 was activated and then neutralized per manufacturer’s protocol. Plates were read using a VERSAmax plate reader (Molecular Devices) and analyzed using MyAssays.com, Microsoft Excel, and GraphPad Prism 9.4.1.

### Cell Sorting

To prepare the highest quality sample for single cell RNA sequencing (scRNA-seq), patients who had the highest percentage of viable, non-neutrophil SFCs were selected, with 3 patients having septic arthritis, and 3 having inflammatory arthritis, regardless of crystal status. As SF in non-pathologic states lacks sufficient cellularity for scRNA-seq analysis, healthy patients were not included. SFCs were thawed in a 37⁰C water bath and diluted in 4 mL Fluorescence Activated Cell Sorting (FACS) buffer and centrifuged at 4⁰C at 1400 rpm. Supernatant was gently decanted, and cells were resuspended in 50 µL of ice cold FACS buffer before staining 1:1000 with DAPI and 5 µl/reaction of: CD244/APC (Clone C1.7), CD11b/BV695 (Clone ICRF44), CD66b/FITC (Clone G10F5) and CD56/PE (Clone 5.1H11) (BioLegend). Cells were incubated in the dark at 4⁰C for 30 minutes, washed once in ice cold FACS buffer, and resuspended in 50 µL of FACS buffer. Samples were then sorted on a Sony MA900 (San Jose, CA) with 100 µm sorting chip, sorting out CD66b^+^ and DAPI positive cells, then sorting on CD11b^+^, CD56^+^, or CD244^+^ positive cells into tubes containing cold PBS with 1% BSA. Cells were then counted for viability using trypan blue on a hemocytometer and concentrated according to 10X Chromium 3’ kit guidelines.

### Single Cell RNA sequencing

Cells were delivered to the Sequencing Core where RNA library generation was performed on a 10X Chromium Controller according to manufacturer’s guidelines. RNA libraries were then sent to NovoGene (Sacramento, CA) for sequencing. Analysis was performed in RStudio with R v4.2.2 using Seurat (11), ReactomeGSA (12) and EnhancedVolcano (13). MT-DNA percentage was limited to 15% during quality control analysis (14). The number of unique genes was set as 200 to 5000. A minimum of 3 cells were required to express each gene. Data was normalized using a global-scaling normalization method per Seurat with a scale factor of 10,000 with log transformation, then the data was integrated into a single database. A resolution of 0.4 was used to define clusters with dimensions set 1:30. Clusters were annotated by identifying top 10 gene expression in addition to expression of markers such as *CD56, CD206, F11R*, and *CD68*. Log2FC threshold was set to 0.5 for gene expression. Analysis was performed comparing septic arthritis to inflammatory arthritis. Adjusted p-values were used to determine significance of Differentially Expressed Genes (DEGs), with Log2FC threshold of 1.5 given the homogenous sample. For Conserved Markers, included genes had min.diff.pct set to 0.7, and min.pct to 0.25.

### Statistics

Descriptive statistics were used to compare patient demographics and underlying diagnoses. Statistical analysis was performed using GraphPad Prism v9.4.1. Flow Cytometry data was tabulated in FlowJo, and 2-way ANOVA with Tukey’s *post-hoc* correction was used to compare %parent or %total cells of control, inflammatory, and septic arthritis populations. Mann-Whitney U-tests were used to compare ELISA results between the 3 arthritis groups (control, inflammatory, and septic). DEG and Conserved gene analysis was performed in Seurat per individual cell cluster using adjusted p-values.

## Results

### Patient Demographics

To identify RSM in SF, 52 patients were recruited into the initial study. Of these, 32% of patients were female and 87% were Caucasian (**Suppl. Table 2**). SF was obtained from 36 of 52 patients. Eighteen patients were found to have septic arthritis based on SF analysis and final culture results, and 8 had inflammatory arthritis. Of the remainder, 10 were designated non-inflammatory (<2000 WBCs) or normal (<200 WBCs) (**Suppl. Table 3**).

### Identification of SF Macrophages Consistent with RSM

We identified a cell subset expressing myeloid marker CD11b, pan-macrophage marker CD68, anti-inflammatory macrophage marker TREM2 (15), Major Histocompatibility Complex marker Class II HLA-DR (16), CX3CR1 (7), OPG (16), and hematopoietic/macrophage adhesion marker CD45 (17). These cells were also negative for Receptor Activator of NF-kB Ligand/RANKL (16) and for the dendritic cell (DC) marker CD11c. There were two distinct subpopulations: a CD14^hi^ and a CD14^dim^ (**Figure 1A**). The CD45^+^CD14^dim^ RSM-like population was found to have an absolute decrease in frequency in pathologic states compared to control SFCs (**Figure 1B**). However, it was also relatively enriched in SF compared to the PBMC fraction (**Figure 1C**). The Alive/CD45^+^CD14^dim^ was then back-gated to better describe this population, first to evaluate the frequency of macrophages by CD68 and TREM2, then if macrophages also co-expressed RSM markers OPG and ZO-1 (**Figure 1D**). For patients with inflammatory arthritis, 1.96% of the CD14^dim^ cells were macrophages, and of those macrophages, 34.35% were RSM-like by OPG and ZO-1 expression. For patients with septic arthritis, 0.92% of CD14^dim^ cells were macrophages, and of those 9.97% were RSM-like.

**Figure 1:**
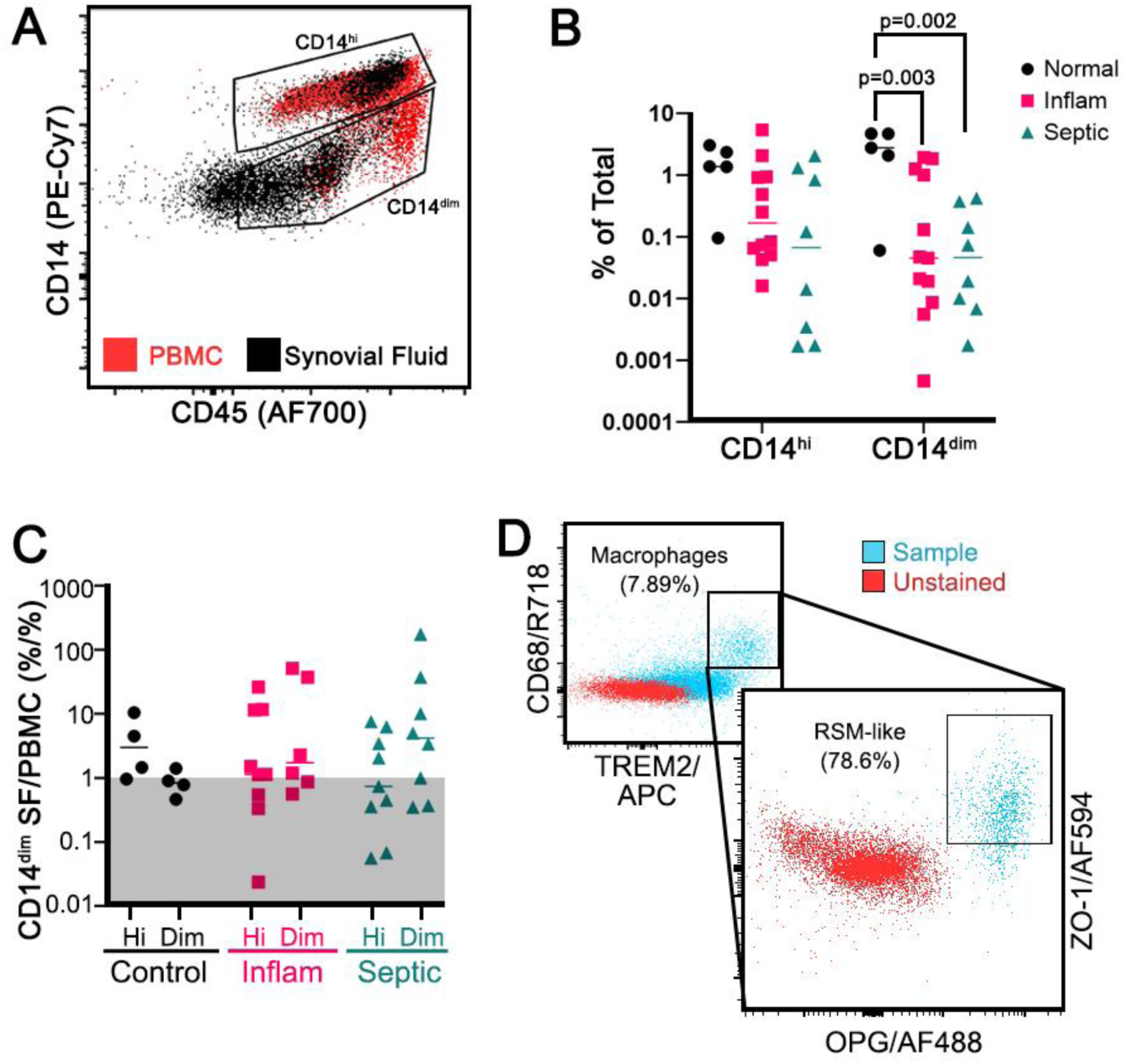
Resident Synovial Macrophage-like cells. Live, CD56^-^CD3^-^CD20^-^CD11c^-^ TREM2^+^OPG^+^CD68^+^CD11b^+^HLA-DR^+^CX3CR1^+^ cells were identified with an enrichment of CD14^dim^ cells in the SF compared to PBMCs (**A**, n=6 patients for preliminary evaluation for RSM cells). These CD14^dim^ macrophages were decreased in absolute frequency in inflammatory and septic arthritis compared to controls (**B**, n=22 patients, 5-12 patients per group) but were relatively enriched in the SF of pathologic joints when SFCs were compared to PBMC, shown by ratio >1 (**C**, n=22 patients, 5-12 patients per group). Back-gating on the Alive/CD14^dim^ population to identify M2 macrophages (D, inset), 78.6% were double positive for OPG and ZO-1 (**D**, representative patient with inflammatory arthritis). Two-way ANOVA with Tukey’s *post-hoc* correction.

### Identification of RSM-like Cells Migrating from Intact Synovium

We obtained synovium from patients undergoing total joint replacement or resection arthroplasty and placed tissue into transwells (**Figure 2A**). This model was validated out 24 hours to have a dose-dependent loss of intimal lining cells with MCP-1 and LPS stimulation (**Suppl. Figure 3**), and negligible apoptosis by Caspace-3 IHC at day 3 in standard culture conditions (**Suppl. Figure 4**). At 24 hours, the migratory synovial myeloid population (Live/CD56^-^/CD3^-^/CD11c^-^/CD20^-^/CD14^+^/CD11b^+^) in a representative patient sample contained OPG+, TREM-2+, ZO-1+, CX3CR1+, and F11R+ populations in the lower transwell media (**Figure 2B**). Sixty-five percent of CD11b+CD14^dim^ cells migrating out of the synovium tissue were double positive for OPG and CX3CR1, and of those cells, 93.8% were double positive for tight junction markers F11R and ZO-1. However, we noted the CD14^hi^ macrophages displayed the greatest increase in the proposed RSM markers (**Figure 2B**), and the relevance of CD14 dim versus high in the acute *ex vivo* setting requires further clarification. To evaluate co-expression of these RSM-specific markers, t-distributed Stochastic Neighbor Embedding (tSNE) plots were created. Of the synovial myeloid population described above, there was weak or low expression of all RSM-specific markers, but ZO-1 may be most specific to identifying migratory RSM in SF (**Figure 2C, top right**). Cells with markers of circulating immune subsets, including neutrophils (CD66b), NK cells (CD56), and T cells (CD3) were also present in the lower transwell chambers (**Suppl. Figure 2**). Though there were no significant differences found between treatments in this experiment, in all cases the stimuli resulted in decreased RSM-like cell migration compared to control, indicating these RSMs may remain active in the tissue, and migratory cells present in the control may be in response to tissue trauma, an effect countered by the stimuli. Cells found adhered to the underside of the transwell or in the bottom of the lower well were below the limit of detection by Crystal Violet assay (data not shown). Therefore, we conclude that RSMs can migrate out of the synovium, and ZO-1 is likely the most specific RSM marker expressed by myeloid cells, but tight junction markers in conjunction with M2 markers and OPG are necessary for identification.

**Figure 2:**
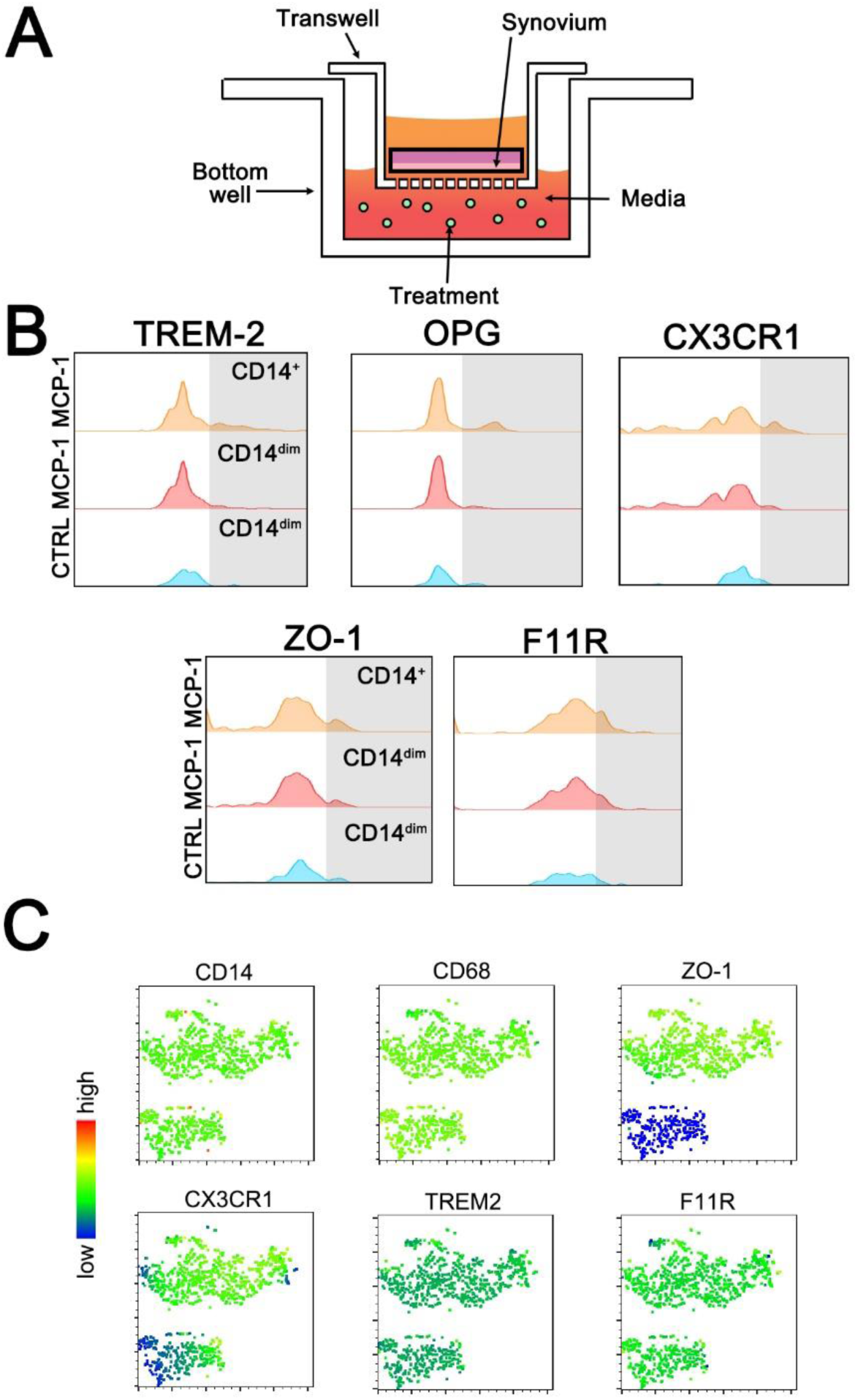
Transwell synovial cell migration assay. Schematic of the experimental set up (**A**). Resident synovial macrophage markers TREM-2+, OPG+, CX3CR1+, ZO-1+ and F11R+ were evaluated within the migratory monocyte/macrophage population (Alive/CD56-/CD3-/CD11c-/CD20-/CD14+/CD11b+) in a representative patient treated with 250 ng/mL of MCP-1 (**B**). tSNE plots of the same patient’s migratory myeloid cells demonstrating ZO-1 has the most delineation from other markers (**C**).

### Evaluation of Pro and Anti-Resorptive Potential of SF

SF supernatant was analyzed for cytokines implicated in bone and/or cartilage destruction. OPG is the decoy receptor for RANKL, which in turn is required for the formation of osteoclasts; the ratio between these two cytokines is an important method to evaluate bone maintenance versus destruction (**Figure 3A**) (18). The collagenase Tartrate Resistant Acid Phosphatase (TRAP), which is secreted by osteoclasts, had no observed change. Transforming Growth Factor Beta (TGF-β) not only inhibits osteoclastogenesis (19), but also stimulates osteoblasts (20) and further, it is stored in the latent phase within the extracellular matrix to be released during bone turnover (21; 22). Likewise, there was no significant difference between pathologic SF and control TGF-β levels (**Figure 3B**). We also tested Bone Morphogenic Protein 2 (BMP2), TNF-α and IFN-γ, however all were below the limit of detection (data not shown). Finally, Tissue Inhibitor of Matrix Metalloproteinases 1 (TIMP1) and Matrix Metallopeptidase 9 (MMP9) were evaluated (**Figure 3C**). TIMP1 is an inhibitor of MMPs. MMP9 is expressed by osteoclasts and is an important enzyme for bone remodeling (23). There was an increase in MMP9 in patients with inflammatory and septic arthritis, and a significantly decreased ratio of TIMP1:MMP9 in these patients as well. Therefore, there is increased potential for bone and cartilage damage in patients with inflammatory and septic arthritis due mainly to increased MMP9, which is possibly due to increased osteoclastogenesis secondary to increased RANKL, or increased synovial fibroblast production. As RSM-like cells were decreased in SF of septic arthritis patients that also had highest RANKL and MMP9, it is possible RSM provide a protective mechanism against bone and joint destruction. The specific cytokine production profile of RSM specifically will require further evaluation.

**Figure 3:**
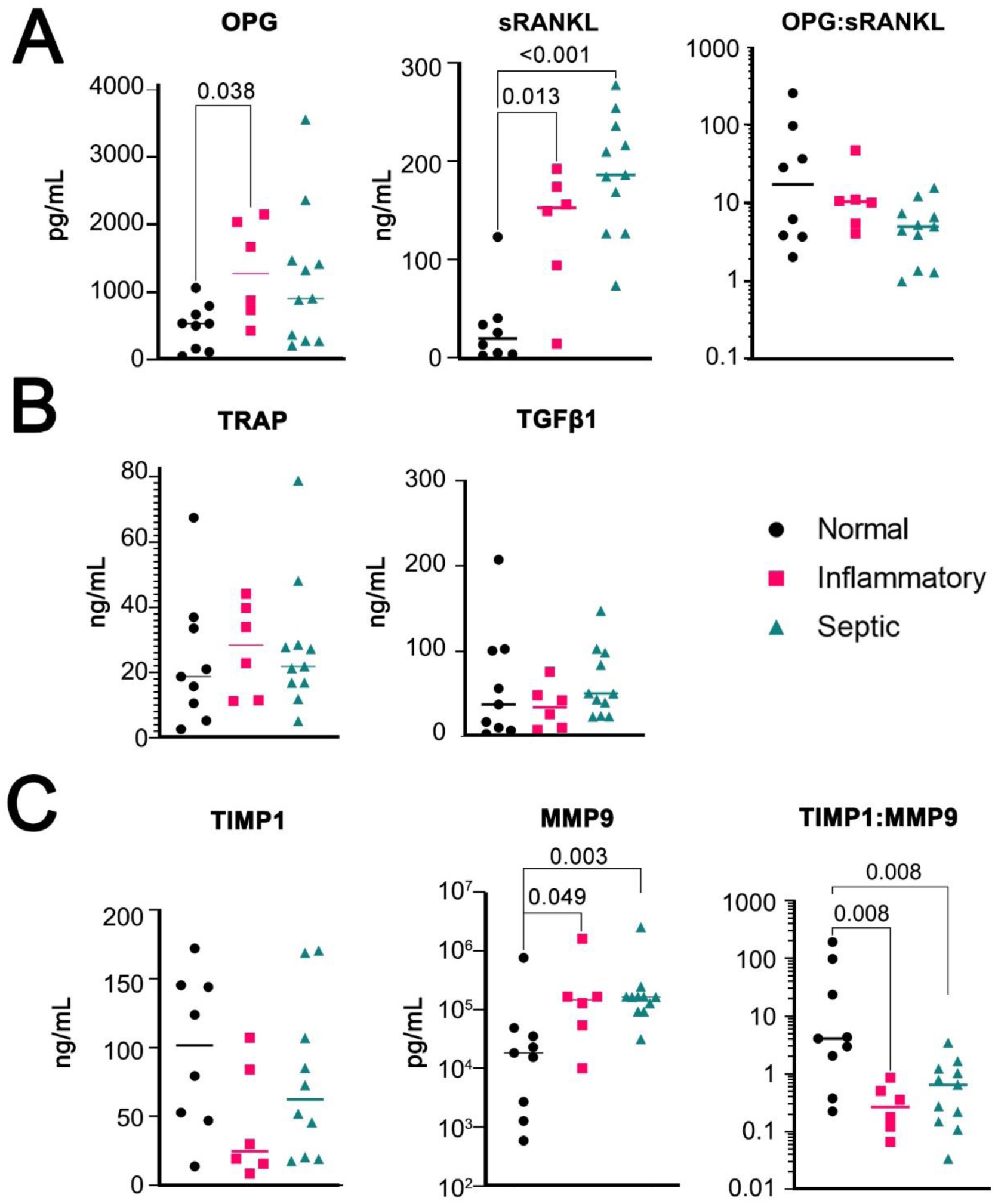
ELISA results of SF supernatant protein concentration. OPG, sRANKL, and ratio of OPG:sRANKL (A). TRAP and TGF-β1, (B). Measures of TIMP1, MMP9, or ratio of TIMP1:MMP9. Normal (n=9), inflammatory (n=6), and septic (n=11). Multiple Mann-Whitney tests with FDR rate <0.05.

### Identification of Rare Cell Subsets Using scRNA-seq

We analyzed sorted myeloid cells from patients with infectious and inflammatory arthritis for highly variable features, demonstrating a relevant focus of M1 and M2 functions (**Suppl. Figure 6A**). Cell subpopulations were clustered into 11 groups (**Figure 4A**) with manual annotation of the clusters based on top ten gene expression (**Figure 4B**). There were insufficient cell events to separately cluster NK and NKT cells, therefore they are represented in a single, though spatially separate, cluster.

**Figure 4:**
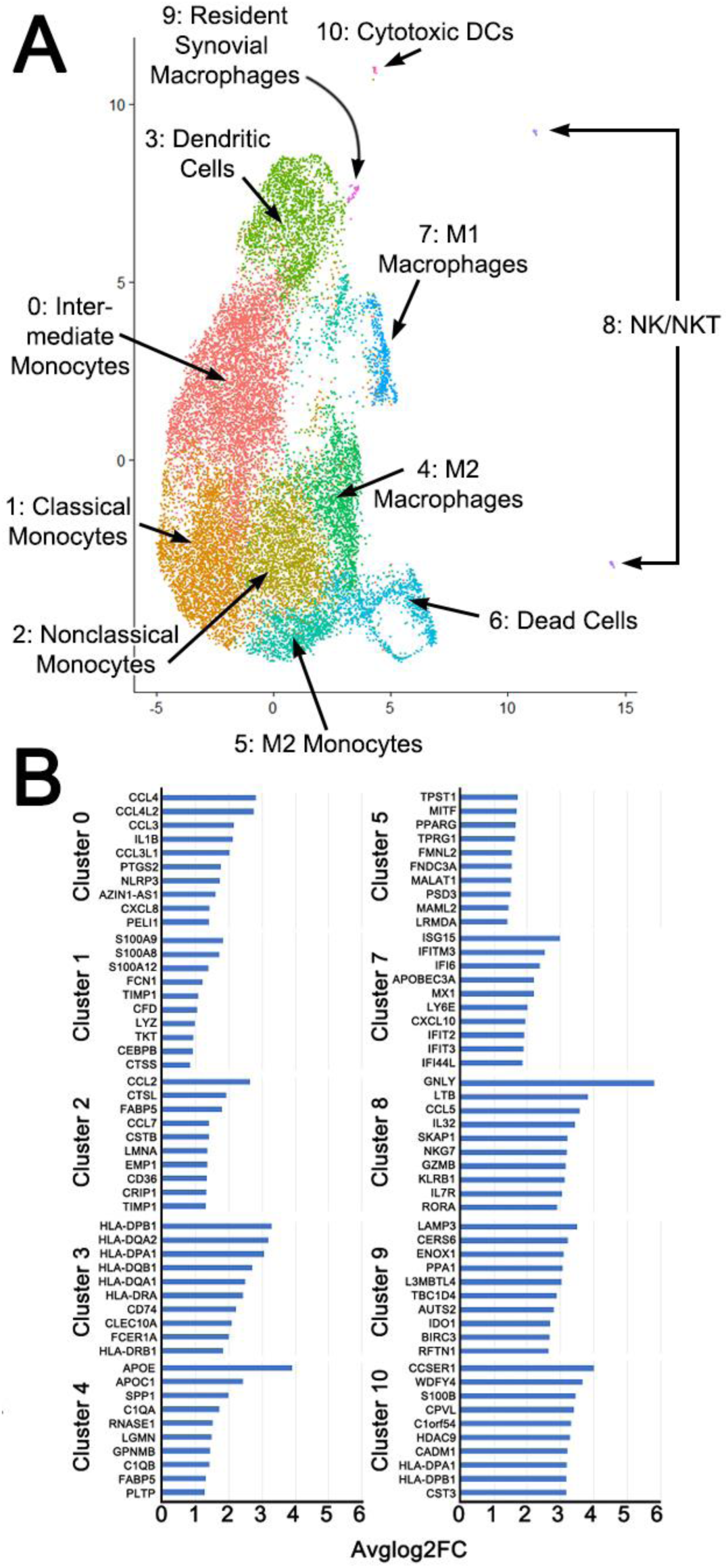
Unsupervised Cluster Analysis showing composite of 3 patients with inflammatory arthritis (A). Clusters were manually identified based on top ten expressed genes (B) in addition to classical markers. Analysis performed in R with resolution of 0.4 and dimensions 1:30. N = 3 patients per inflammatory and septic arthritis.

After assigning known and widely accepted monocyte/macrophage designations based on gene profiles, clusters 9 and 10 remained. Cluster 10 expressed NK-marker *CD56,* low *CD68* (**Suppl. Figure 6B**), while also having high *HLA* expression (**Figure 4B**). However, Cluster 10 did not express any granzymes. Therefore, we putatively classified this cluster as cytotoxic DCs (24). Cluster 9 expressed *F11R* (7), *CD68,* and M2-marker *IDO1* (25) (**Figure 4B, Suppl. Figure 6B**). It was also the only cluster with *OPG* expression, though this was not significant. This is putatively consistent with the RSM phenotype. *ZO-1* was not expressed in any cluster. We also evaluated expression of resident macrophage transcription factor *GATA6* (26), which was minimally expressed, but only in Cluster 3. This may indicate that RSMs are split between multiple clusters, as Cluster 3 was designated as DCs based on high expression of HLA—which is also consistent with sub-intimal RSM, but not intimal RSM (27). Based on PCA analysis (**Suppl. Figure 6C**), cytotoxic DCs were more closely related to NK/NKT cells than to RSM, and the cytotoxic DC and RSM populations may represent phases of differentiation of the same cell of origin, as both were distinct from the circulating monocyte/macrophage population. The plasticity of tissue macrophages, and ability to survive in new compartments has been advocated and challenged; the exact lineage remains unknown.

Differentially Expressed Genes (DEGs) were explored using a volcano plot (**Suppl. Figure 6D**) and a Log2FC threshold of 1.5. DEGs were statistically significant in Clusters 0-8 (**Figure 5A**). Transcripts highly upregulated in septic arthritis (and therefore down-regulated in inflammatory arthritis) among multiple clusters included *HLA-DRB5, C15orf48, IL1B, AC025580.2*, and *SOD2*. In inflammatory arthritis, common gene upregulation included *PPARG, FABP5, CD36*, *NLRP3, SPP1, and MITF*. M1 macrophages from the two different arthritis etiologies showed a distinctly different gene expression patterns with an upregulation of *GBP1, GBP5*, and *TIMP1*.

**Figure 5:**
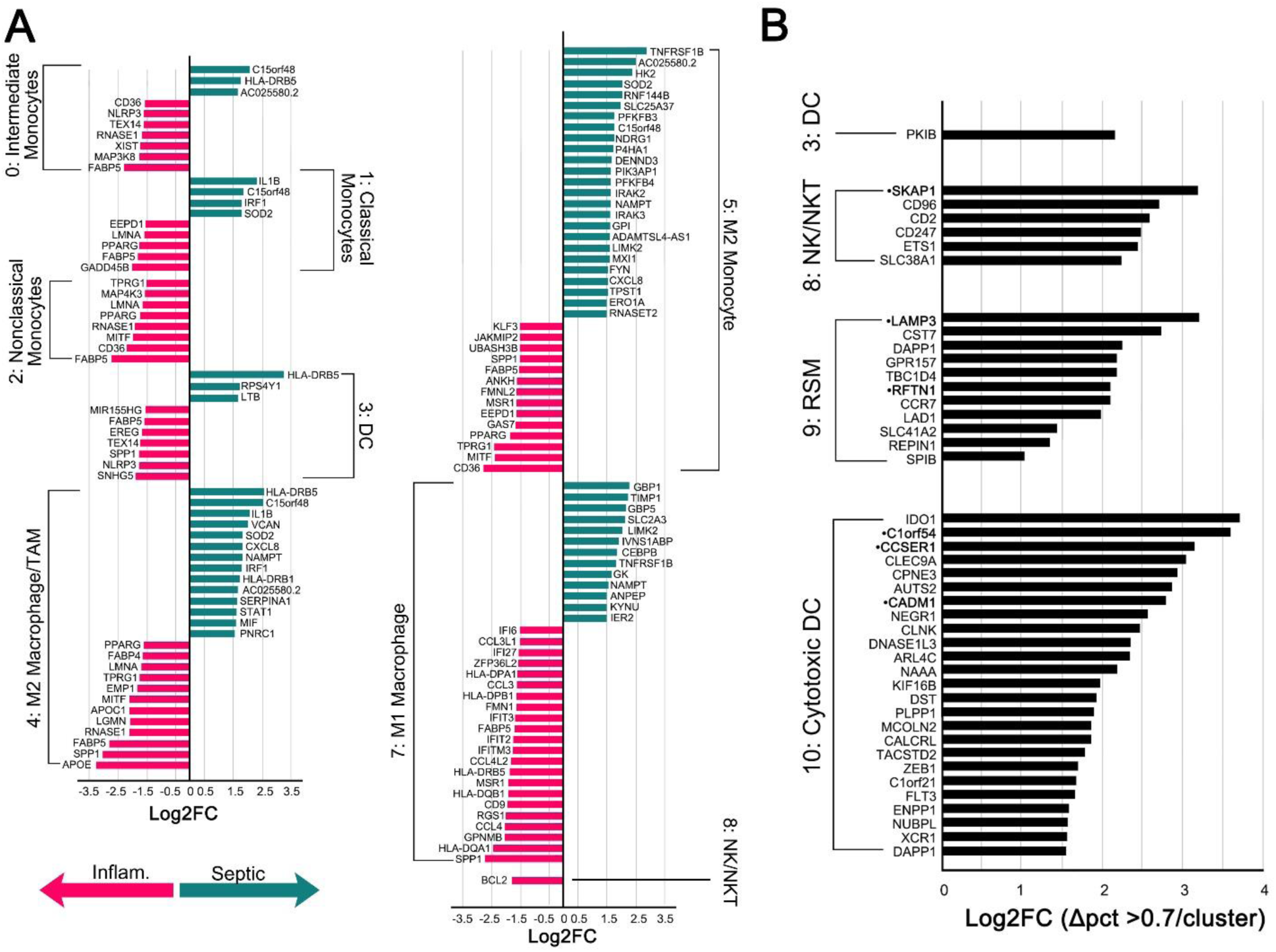
Differentially Expressed Genes (DEGs) (**A**). Conserved genes where Log2FC > 1.5 and the change in percent expression in both septic arthritis and inflammatory arthritis was > 0.7 in each cluster compared to all other clusters. Gene names in **•bold** represent genes that were also in the top 10 expressed genes in Figure 4.

Conserved genes that remained highly expressed in both inflammatory and infectious arthritis were also examined, as these could represent novel targets in the treatment, or prevent the conversion of infectious to inflammatory arthritis. Conserved genes were only significant in DC, NK/NKT, RSM, and Cytotoxic DC clusters (**Figure 5B, Suppl. Table 4**). Nearly all genes were associated with cytolytic function, antigen presentation, M1/M2 polarity, or lysosomes.

Gene Set Analysis (GSA) was then assessed to determine the overall, broad picture, of cell function and metabolism (**Figure 6**) with the 15 most upregulated or downregulated pathways, which identified complement, bone-derived FGF23 signaling, and COX signaling. Pathways specifically immune relevant, related to phagocytic potential, complement associated and adhesion related (**Suppl. Figure 7**) were also examined. In all cases except adhesion, Cytotoxic DC, NK/NKT, and RSMs populations shared a similar pattern of activity, indicating functional overlap. GSA was further performed specifically on the RSM cluster to identify the maximum changes in septic and inflammatory arthritis (**Suppl. Figure 8**), which identified threonine, pyridoxine, and thiamine metabolism.

**Figure 6:**
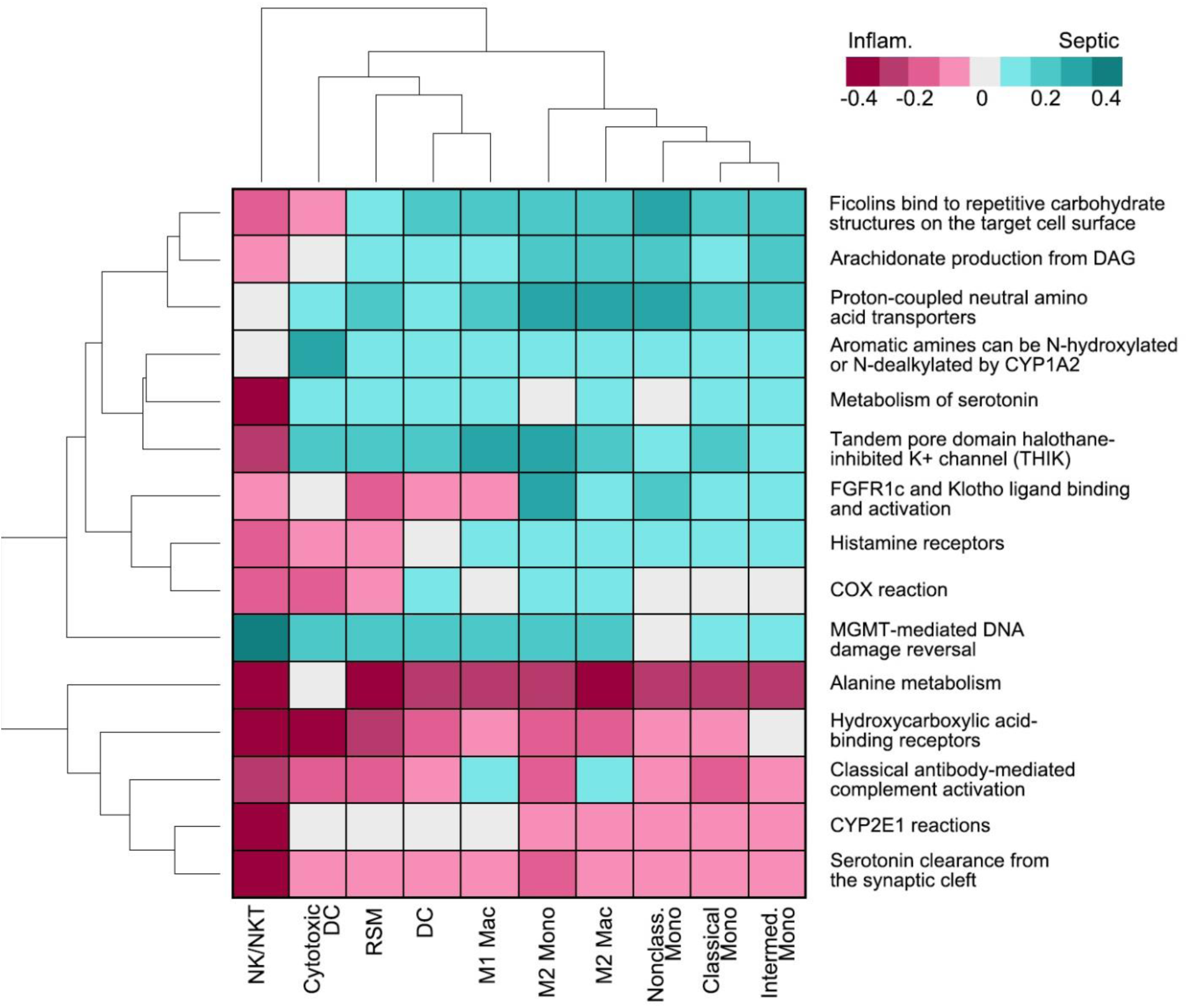
Top 15 most up or downregulated pathways using DEGs and ReactomeGSA.

## Discussion

RSMs are regulatory immune cells of the joint space that are historically identified using tissue histology. We identified a rare subset of SF macrophages identified by flow cytometry consistent with previously described RSMs, indicating migratory capacity especially during pathologies that dysregulate tight junctions. The role of these cells in SF has yet to be established.

Though the chemokine receptor CX3CR1 was found to be specific for murine RSM (1), we were unable to distinguish SFCs from peripheral monocytes using this marker. Likewise, HLA-DR expression was downregulated in the intimal-lining RSMs, but upregulated in sub-lining RSMs (8), and it is unclear if HLA-DR expression would assist in identification of RSM-like cells in the SF. We first identified putative RSMs as CD14^dim^OPG^+^ M2-macrophages, and while CD14^dim^ monocytes have been described previously as non-classical and poorly phagocytic (28; 29), data is limited on CD14^dim^ macrophages. CD14^dim^ gingival macrophages were M2 and likely osteo-protective by high expression of IL-10 and TGF-β in the setting of gingivitis (30). To fully elucidate the utility of CX3CR1, HLA-DR, and CD14 in human SF macrophage subsets requires further work.

To determine whether our identified cells came from synovium or from circulation, we piloted a novel transwell migration assay with human *ex vivo* synovium that validated our flow cytometry findings. Therefore, we believe the CD68/TREM2/OPG/ZO-1/F11R myeloid cell fraction is the most representative of putative RSMs. This explant model could be used widely to study other synovial pathology.

We then utilized scRNA-seq to identify rare macrophage subpopulations. Markers identified using flow cytometry were not always identified in gene transcripts, including *ZO-1.* Instead we identified M2/Tumor Associated Macrophage markers such as *IRF4* (31) and *IDO1* (32) (**Suppl. Figure 6B**). While the putative RSM cluster resembled M2 macrophages, these cells also had a similar transcriptional signature to inflammatory NK/NKT and Cytotoxic DCs, indicating that RSMs may be capable of taking on an inflammatory and/or joint destructive phenotype in settings of chronic inflammation. This was demonstrated by the pro-inflammatory expression of *CCR7* (Log2FC 2.25) and *CD86* (Log2FC 0.92) in infectious settings.

To provide context to our findings, we compared our findings to scRNA-seq performed on synovium by other groups. Human MERTK^+^CD206^+^ RSMs were anti-inflammatory in patients in remission from rheumatoid arthritis (RA) (27). We found *MERTK* expressed highest in Cluster 5/M2 Monocytes (Log2FC 0.94), but it was not expressed in 9/RSM. Interestingly, MERTK^-^CD206^-^ RSM indicated active RA (27). As all patients who received arthrocentesis and participated in the scRNA-seq were acutely symptomatic, a MERTK^+^CD206^+^ RSM profile was likely physiologically improbable. For *CD206,* this was expressed highest in Cluster 4/M2 Macrophages, which additionally expressed *TREM2, FOLR2*, and *LYVE1* (27), though only *TREM2* had Log2FC >0.5. Once RSMs exit synovial tissue to enter the SF, previously established profiles may no longer apply, and RSMs may be distributed amongst clusters rather than a discrete cluster.

Others found that RSMs arose from *CSF1R+* interstitial macrophages (7). We found *CSF1R* expressed in clusters 3-6, and 8-10. It was also found that interstitial RSM expressed *RETNLA, STMN1*, and *AQP1* (7). *RETNLA* and *AQP1* transcripts were not identified in this study, while *STMN1* was only expressed by Cluster 3/DCs, but this was not significant. Likewise, markers of tight junctions and cell polarity previously found included *F11R, CLDN5, FAT4*, and *VANGL2*. Of these, only *F11R* was identified, and expressed mainly in RSM and Cytotoxic DC clusters (**Suppl. Figure 6B**).

We identified *LAMP3/CD208/DC-LAMP* as the top gene expressed by Cluster 9/RSM, and while the exact function of LAMP3 has yet to be elucidated, it is likely to be involved with MHC Class II peptide presentation (34). *LAMP3* is also traditionally considered a marker of mature DCs. DCs expressing LAMP3 are regulatory in nature and more enriched in draining lymph nodes rather than tumors (35). However, *LAMP3* is upregulated by THP-1 macrophages with *in vitro* LPS stimulation (36), and is constitutively expressed in primary macrophages in multiple species (37). Further analysis of Cluster 9/RSM gene expression revealed high expression of *ENOX1* (**Figure 4**), involved with reduction of oxygen to superoxide (38). Likewise, *CERS6* contributes to mitochondrial dysfunction by promoting reactive oxygen species production in hepatocytes (39). *IDO1* expression in macrophages has been associated with increased tumor immune cell infiltration (40) and tryptophan metabolism, the metabolites of which inhibit oxidative cell death (41). Together this indicates that these cells are likely to have potent generation of superoxide with inhibition of apoptosis, which may indicate perpetuation of chronic inflammation. Until these putative RSM can be compared to similar cells from healthy SF, the baseline role and function are unclear.

Given we believe these cells are capable of migratory function, markers of migration and extravasation were also examined. *CADM1* was highly and conservatively expressed (**Figure 5**) and is strongly associated with TREM2+ tumor associated macrophages (42). *ALCAM* (Log2FC 0.73) is expressed by endothelial cells of the blood-brain barrier and migrating monocytes (43), which may be consistent with the relative immune privilege of the joint space. As ALCAM stabilizes tight junctions (44), this integrin may be important in the homeostasis of the joint space as maintained by RSM. *PECAM1* (Log2FC -1.03) assists with leukocyte migration through tight junctions (45). Our data seems to suggest a differential regulation of tight junctions by macrophages depending on pathology.

As there were no DEGs identified in Cluster 9/RSM, GSA was performed to identify unique pathways and discovered threonine, pyridoxine, and thiamine metabolism (**Suppl. Figure 8**). The role of threonine catabolism is unclear in macrophages but is critically necessary for murine stem cell viability (46), which may be similar function to locally renewing macrophage populations. Pyridoxine suppresses IL-1β release through NLRP3 inhibition (47), while thiamine precursors have been found to both increase cellular glutathione stores and inhibit NF-kB translocation to the nucleus in microglial cells (48), the resident macrophages of the brain. Putative RSMs are upregulating anti-inflammatory pathways, but the question remains if the concurrent upregulation of complement and superoxide transcripts may supersede the protective mechanisms in place through B vitamins.

We also evaluated the protein levels of multiple bone-relevant cytokines and related this to cell populations in the joint space. The DEG *TIMP1* was identified in Cluster 7/M1 Macrophage as highly upregulated in septic arthritis, with concurrent downregulation in inflammatory arthritis. As a 1:1 inhibitor of MMP9 (49), TIMP1 is likely protective in the setting of joint inflammation by preserving the extracellular matrix of cartilage and bone from enzymatic degradation. We observed a decrease in the stoichiometric ratio of TIMP1:MMP9 protein concentration in both infectious and inflammatory arthritis SF, which is concerning for MMP9 as a major cause of bone destruction in these pathologic states, and a potential therapeutic target. *MMP9* was not differentially expressed, and MMP9 found by ELISA is likely from other cells, specifically fibroblast like synoviocytes and/or neutrophils (50), but possibly also osteoclasts (23). *TIMP1* was expressed in Cluster 9/RSM (Log2FC -2.67) indicating RSM would not be the primary source of this in SF.

In conclusion, the profile of a subset of M2, likely osteoprotective macrophages in SF that can be stimulated to migrate out of synovial tissue *ex vivo* suggests these are equivalent to tissue resident macrophages. However, these putative RSM expressed transcripts heavily involved with antigen presentation (*LAMP3*), oxidative stress (*ENOX1, CERS6, IDO1*), and cell migration (*ALCAM, PECAM*). This suggests that in settings of infectious or inflammatory arthritis, these cells may perpetuate rather than attenuate inflammation, and could be involved in the transition to chronic symptomology once out of the normal synovial tissue niche. Further work will focus on the cytokine expression of these putative migratory RSM, and the interactions of RSM with other cells in the joint space, including T cells, B cells, and NK or NKT cells.

## Acknowledgements

The authors would like to thank Catherine Fairfield BSN, EM Research Coordinator, and Alex Peebles, Cameron Williams, Shannon Landers, Klaudia Golebiewski, Allison Herr, Noble Briggs, Vanko Bicar, Malea Pinckney, Heath Gibbs, Nicole Grossmann, Scott Tibbetts, Ike Appleton, Jay Miller, and Jacob Hampton with the Emergency Department Research Enroller Program (ED-REP) for their essential role in enrolling patients and collecting data for this project.

The authors would also like to thank Dr. Daniel Livorsi and the Division of Infectious Diseases for assistance with patient recruitment.

The data presented herein were obtained at the Flow Cytometry Facility, which is a Carver College of Medicine / Holden Comprehensive Cancer Center core research facility at the University of Iowa. The facility is funded through user fees and the generous financial support of the Carver College of Medicine, Holden Comprehensive Cancer Center, and Iowa City Veteran’s Administration Medical Center. Research reported in this publication was supported by: the National Center for Research Resources of the National Institutes of Health under Award Number 1 S10 OD034193-01; and the National Cancer Institute of the National Institutes of Health under Award Number P30CA086862

---

**Supplementary Table 1:** Flow cytometry panel, initial (top) and modified (bottom). Mac = macrophage; Mono = monocyte; OB = osteoblast; RSM = Resident Synovial Macrophage; APC = Antigen Presenting Cell

| Molecular Target | Fluorochrome | Clone | Vendor | Cell of Interest |
| --- | --- | --- | --- | --- |
| CD11b | BV605 | ICRF44 | BioLegend | Myeloid |
| CD11c | BV785 | 3.9 | BioLegend | Dendritic Cell |
| CD14 | PE-Cy7 | 63D3 | BioLegend | Mac/Mono |
| CD20 | BV785 | 2H7 | BioLegend | B cell |
| CD3 | BV650 | UCHT1 | BioLegend | T Cell |
| CD45 | AF700 | 2D1 | BioLegend | Leukocytes |
| CD56 | PE-Cy5 | 5.1H11 | BioLegend | NK cell |
| CD68 | PerCP-Cy5.5 | Y1/82A | BioLegend | Macrophage |
| CX3CR1 | BV421 | 2A9-1 | BioLegend | RSM/Mac/Mono |
| HLA-DR | PE | Tu36 | BioLegend | Activated APC |
| OPG/TNFRSF11B | AF488 | 155321 | R&D Systems | RSM |
| RANKL/TNFSF11 | AF350 | 685857 | R&D Systems | OB/activated T cells |
| TREM2 | APC | 237920 | R&D Systems | Macrophage |
| Viability | Live/Dead Aqua |  | Thermofisher |  |
| CD11b | BV605 | ICRF44 | BioLegend | Myeloid |
| CD11c | BV785 | 3.9 | BD Biosciences | Dendritic Cell |
| CD14 | Spark Blue 550 | 63D3 | BioLegend | Mac/Mono |
| CD20 | BV785 | 2H7 | BD Biosciences | B cell |
| CD3 | BV510 | SK7 | BioLegend | T Cell |
| CD45 | AF700 | HI30 | BioLegend | Leukocytes |
| CD56 | PE-Cy5 | 5.1H11 | BioLegend | NK cell |
| CD66b | PE | MIH24 | BioLegend | Neutrophil |
| CD68 | R718 | Y1/82A | BD Biosciences | Macrophage |
| CX3CR1 | BV711 | 2A9-1 | BioLegend | RSM/Mac/Mono |
| F11R/JAM-1/JAM-A | BV421 | M.Ab.F11 | BD Biosciences | RSM |
| OPG/TNFRSF11B | AF488 | 155321 | R&D Systems | RSM |
| TREM2 | APC | 237920 | R&D Systems | Macrophage |
| ZO-1/TJP1 | Coralite 594 | polyclonal | ThermoFisher | RSM |
| Viability | Live/Dead Blue |  | ThermoFisher |  |

**Supplemental Table 2:** Patient Demography. Arthritis Type was determined by synovial fluid WBC count per mm^3^, %PMNs, and/or culture results. * = 3 patients had Lyme Arthritis, 1 of which had both a *S. aureus* and positive IgG with confirmatory Western Blot.

| Demographics n=52 |  |  |
| --- | --- | --- |
| **Age** (years) |  |  |
|  | Mean (STD) | 51.7 (21) |
|  | Median | 58 |
|  | Range | 3-77 |
| **Gender** n, (%) |  |  |
|  | Male | 35 (68) |
|  | Female | 17 (32) |
| **Race and Ethnicity** n, (%); Hispanic n, (%) |  |  |
|  | White | 45 (87); 1 (2) |
|  | Black | 3 (6); 0 |
|  | Asian | 1 (2); 0 |
|  | Multiracial | 1 (2); 0 |
|  | Unknown | 1 (2); 0 |
| **Arthritis Type** n (%) |  |  |
|  | Normal/Non-I | 9 (25) |
|  | Inflammatory | 8 (22) |
|  | Septic* | 18 (50) |
|  | Hemorrhagic | 1 (3) |

**Supplementary Table 3:** Patient Diagnosis. SF analysis was performed on patients when it was clinically indicated. SF diagnosis was based on WBC count (# nucleated cells) and % PMN based on neutrophil count. * = not performed. ** = Lyme Arthritis in addition to other bacterial arthritis.

| Patient | Color | Clarity | Crystals | # Nucleated Cells | # Neutrophils | # Lymphocytes | Arthritis Type |
| --- | --- | --- | --- | --- | --- | --- | --- |
| 2 | Red | Turbid | CCP, MSU | 29783 | 22635 | 101 | Inflammatory |
| 3 | Yellow | Turbid | CPP | 4716 | 2453 | 20 | Inflammatory |
| 5 | Red | Turbid | none | 822 | 674 | 107 | Non-Inflammatory |
| 6 | Yellow | Turbid | none | 29497 | 28022 | 0 | Septic |
| 7 | Yellow | Hazy | none | 714 | 29 | 64 | Non-Inflammatory |
| 9 | Yellow | Hazy | none | 63726 | 60539 | 1275 | Septic |
| 10 | Yellow | Slightly Hazy | none | 464 | 28 | 70 | Non-Inflammatory |
| 11 | Red | Turbid | none | 671 | 658 | 0 | Non-Inflammatory |
| 13 | Orange | Hazy | none | 831 | 632 | 58 | Non-Inflammatory |
| 14 | None | Clear | none | 29 | 0 | 7 | Normal |
| 15 | Orange | Turbid | none | 63184 | 60025 | 632 | Inflammatory |
| 16 | Pink | Hazy | none | 166627 | 128303 | 9998 | Septic |
| 18 | Amber | Turbid | none | 134280 | 119509 | 2686 | Septic |
| 19 | Red | Turbid | none | 11522 | 8987 | 1959 | Inflammatory |
| 21 | N/A | N/A | none | N/A | N/A | N/A | Septic |
| 22 | Yellow | Turbid | CPP | 34482 | 27586 | 2 | Inflammatory |
| 23 | Yellow | Clear | none | 348 | 28 | 52 | Non-Inflammatory |
| 24 | Red | Turbid | none | 1819 | 1564 | 14 | Hemorrhagic |
| 26 | Yellow | Clear | none | 26 | 5 | 12 | Normal |
| 27 | Orange | Turbid | MSU | 33805 | 26030 | 0 | Inflammatory |
| 28 | Brown | Turbid | none | 66945 | * | * | Septic |
| 30 | Yellow | Slightly Hazy | none | 26536 | 25209 | 531 | Septic |
| 31 | Red | Hazy | none | 48363 | 46428 | 484 | Septic |
| 32 | Orange | Hazy | none | 43355 | 42054 | 434 | Septic |
| 33 | Red | Turbid | none | 304 | 76 | 85 | Septic |
| 34 | Red | Hazy | none | 854 | 384 | 154 | Non-Inflammatory |
| 38 | Red | Turbid | none | 114000 | 1000320 | 2 | Septic |
| 39 | Red | Turbid | MSU | 76890 | 76121 | 0 | Inflammatory |
| 40 | Pale Yellow | Turbid | CPP | 67368 | 64673 | 0 | Inflammatory |
| 41 | Pink | Turbid | None | 79118 | 75162 | 0 | Septic** |
| 43 | Orange | Turbid | None | 104595 | 92044 | 2 | Septic |
| 46 | Pink | Turbid | None | 80562 | 74117 | 2417 | Septic |
| 47 | Red | Turbid | None | 9884 | 8303 | 692 | Septic |
| 48 | Orange | Hazy | None | 130 | 83 | 26 | Normal |
| 49 | Yellow | Hazy | MSU | 20566 | 18715 | 0 | Inflammatory |

**Supplementary Table 4:** List of all significant conserved genes in RSM, Cytotoxic DC, and NK/NKT clusters.

| Gene | Log2FC | $\Delta$ % Septic | $\Delta$ % Inflamm | Cluster | Gene Function |
| --- | --- | --- | --- | --- | --- |
| PKIB | 2.10657 | 0.71 | 0.726 | DC | Also known as AKT, causes M2 formation |
| SKAP1 | 3.16933 | 0.89 | 0.954 | NK/NKT | TCR adaptor protein |
| CD96 | 2.690931 | 0.77 | 0.874 | NK/NKT | Inhibitory, marker of exhaustion |
| CD2 | 2.571958 | 0.78 | 0.817 | NK/NKT | Regulates lytic activity |
| CD247 | 2.473159 | 0.79 | 0.869 | NK/NKT | Subunit of T cell antigen receptor complex |
| ETS1 | 2.430819 | 0.85 | 0.88 | NK/NKT | Required for NK differentiation and cytotoxic function |
| SLC38A1 | 2.222173 | 0.82 | 0.77 | NK/NKT | Glutamine transporter |
| IDO1 | 3.679276 | 0.93 | 0.764 | Cytotoxic DC | Immunosuppressive, expressed by DCs, macs, and epithelial cells |
| C1orf54 | 3.555745 | 0.94 | 0.794 | Cytotoxic DC | cDC1 gene |
| CCSER1 | 3.110627 | 0.78 | 0.818 | Cytotoxic DC | upregulated in anti-inflammatory conditions |
| CLEC9A | 3.009585 | 1.00 | 0.888 | Cytotoxic DC | DC receptor for necrotic cell death |
| CPNE3 | 2.908024 | 0.82 | 0.76 | Cytotoxic DC | cDC1 gene |
| AUTS2 | 2.836438 | 0.78 | 0.74 | Cytotoxic DC | associated with autism |
| CADM1 | 2.766723 | 0.99 | 0.936 | Cytotoxic DC | expressed by pre-cDC1, cell adhesion |
| NEGR1 | 2.54191 | 0.97 | 0.798 | Cytotoxic DC | immunoglobulin superfamily cell adhesion molecule subgroup IgLON, has been implicated in neuronal growth and connectivity |
| CLNK | 2.441584 | 0.99 | 0.885 | Cytotoxic DC | upregulated in response to IL-2 and IL-3, associated with SLP-76 |
| DNASE1L3 | 2.326661 | 0.89 | 0.998 | Cytotoxic DC | Cytokine secretion following inflammasome activation in response to DNA in circulating apoptotic bodies |
| ARL4C | 2.315802 | 0.71 | 0.709 | Cytotoxic DC | oncogene, tubulogenesis, wnt-beta catenin signaling |
| NAAA | 2.166039 | 0.71 | 0.78 | Cytotoxic DC | inflammatory, lysosomal |
| KIF16B | 1.944567 | 0.79 | 0.778 | Cytotoxic DC | microtubule formation associated with endosomes and cross presentation |
| DST | 1.91132 | 0.77 | 0.764 | Cytotoxic DC | junctional adhesion protein |
| PLPP1 | 1.872874 | 0.74 | 0.736 | Cytotoxic DC | phospholipid lipase associated with macro inflammation, cDC1 marker |
| MCOLN2 | 1.841604 | 0.73 | 0.824 | Cytotoxic DC | MHCII presentation, innate immune cell activation, enhances infectivity of certain viruses, interferon stimulating gene |
| CALCRL | 1.840268 | 0.76 | 0.951 | Cytotoxic DC | regulates DC function through NF-kB |
| TACSTD2 | 1.765963 | 0.78 | 0.989 | Cytotoxic DC | Marker for TGF-B1 dependent DCs |
| ZEB1 | 1.672937 | 0.74 | 0.941 | Cytotoxic DC | cDC1 cells |
| C1orf21 | 1.647088 | 0.84 | 0.9 | Cytotoxic DC | enriched in cytotoxic CD8 T cells |
| FLT3 | 1.645133 | 0.96 | 0.895 | Cytotoxic DC | The growth factor Flt3 ligand (Flt3L) is central to dendritic cell (DC) homeostasis and development, controlling survival and expansion by binding to Flt3 receptor tyrosine kinase on the surface of DCs. |
| ENPP1 | 1.548458 | 1.00 | 0.777 | Cytotoxic DC | suppresses innate immune response, |
| NUBPL | 1.540362 | 0.83 | 0.911 | Cytotoxic DC | mitochondrial gene, iron and sulfur |
| XCR1 | 1.53506 | 0.78 | 0.995 | Cytotoxic DC | DC, induces CD8 responses |
| DAPP1 | 1.52997 | 0.72 | 0.717 | Cytotoxic DC | immune related mucosal tissues, hyper-reactive airway disease, IDO1/CCR7 signaling, associated with DCs |
| LAMP3 | 3.181144 | 0.88 | 0.967 | RSM | lysosome |
| CST7 | 2.710421 | 0.93 | 0.748 | RSM | Cistatin F, lysosomal cathepsin inhibitor |
| DAPP1 | 2.225168 | 0.72 | 0.781 | RSM | immune related mucosal tissues, hyper-reactive airway disease, IDO1/CCR7 signaling, associated with DCs |
| GPR157 | 2.163834 | 0.75 | 0.897 | RSM | G protein coupled receptor 157 |
| TBC1D4 | 2.161417 | 0.79 | 0.941 | RSM | Rab GTPase Activating Protein |
| RFTN1 | 2.085313 | 0.77 | 0.736 | RSM | glucose transport |
| CCR7 | 2.083676 | 0.92 | 0.837 | RSM | M1 polarization |
| LAD1 | 1.96187 | 0.77 | 0.713 | RSM | epithelialization |
| SLC41A2 | 1.415962 | 0.76 | 0.78 | RSM | magnesium transporter |
| REPIN1 | 1.325367 | 0.76 | 0.72 | RSM | increasing cell size and tumor metastasis |
| SPIB | 1.005641 | 0.75 | 0.726 | RSM | recruits TAMs via CCL4 signaling |

**Supplemental Figure 1:**
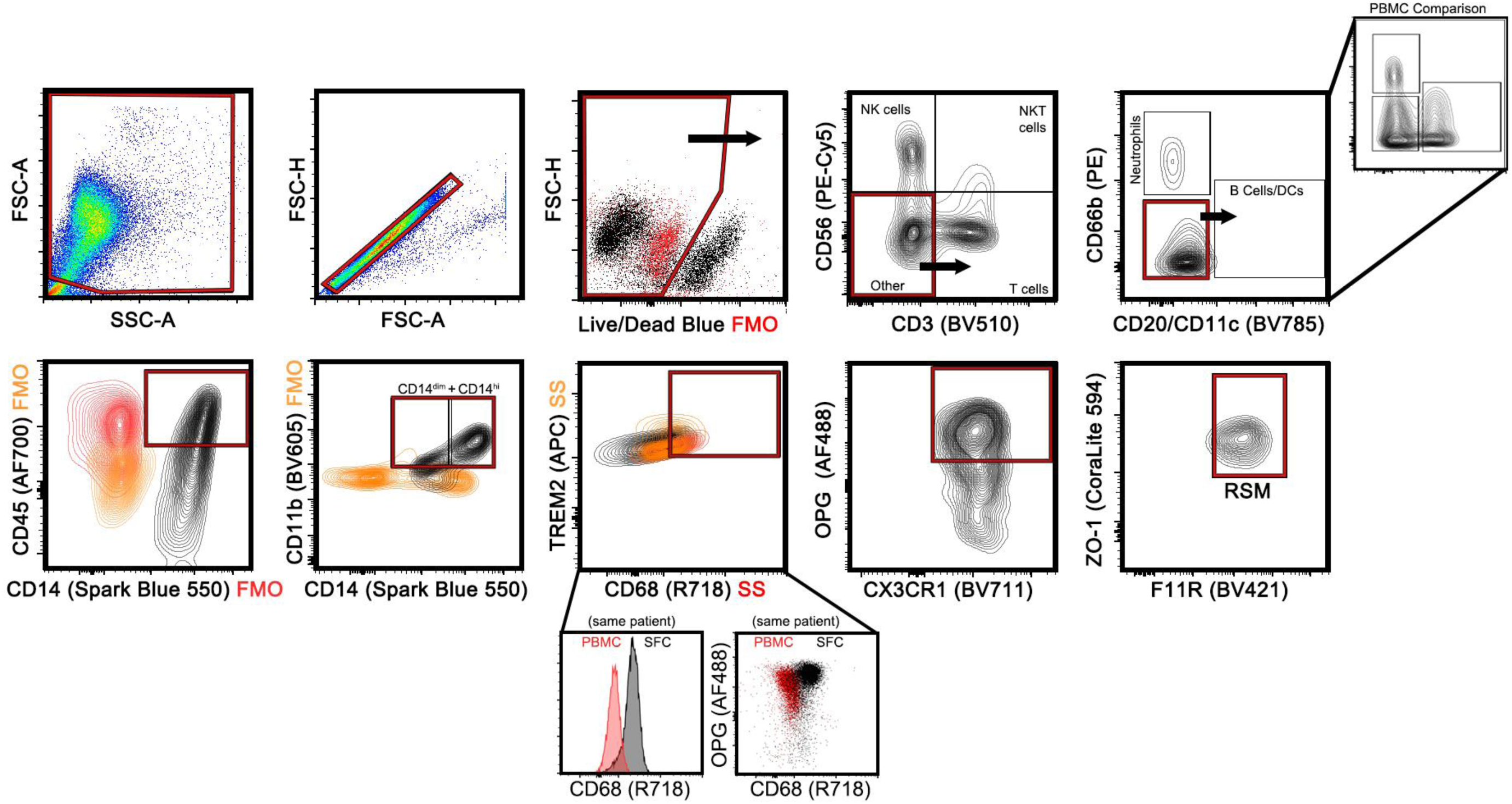
Final gating strategy to identify TREM2^+^CX3CR1^+^OPG^+^F11R^+^ZO-1^+^ macrophages consistent with RSM. FMO and single stain (SS) shown where used to define gates.

**Supplemental Figure 2:**
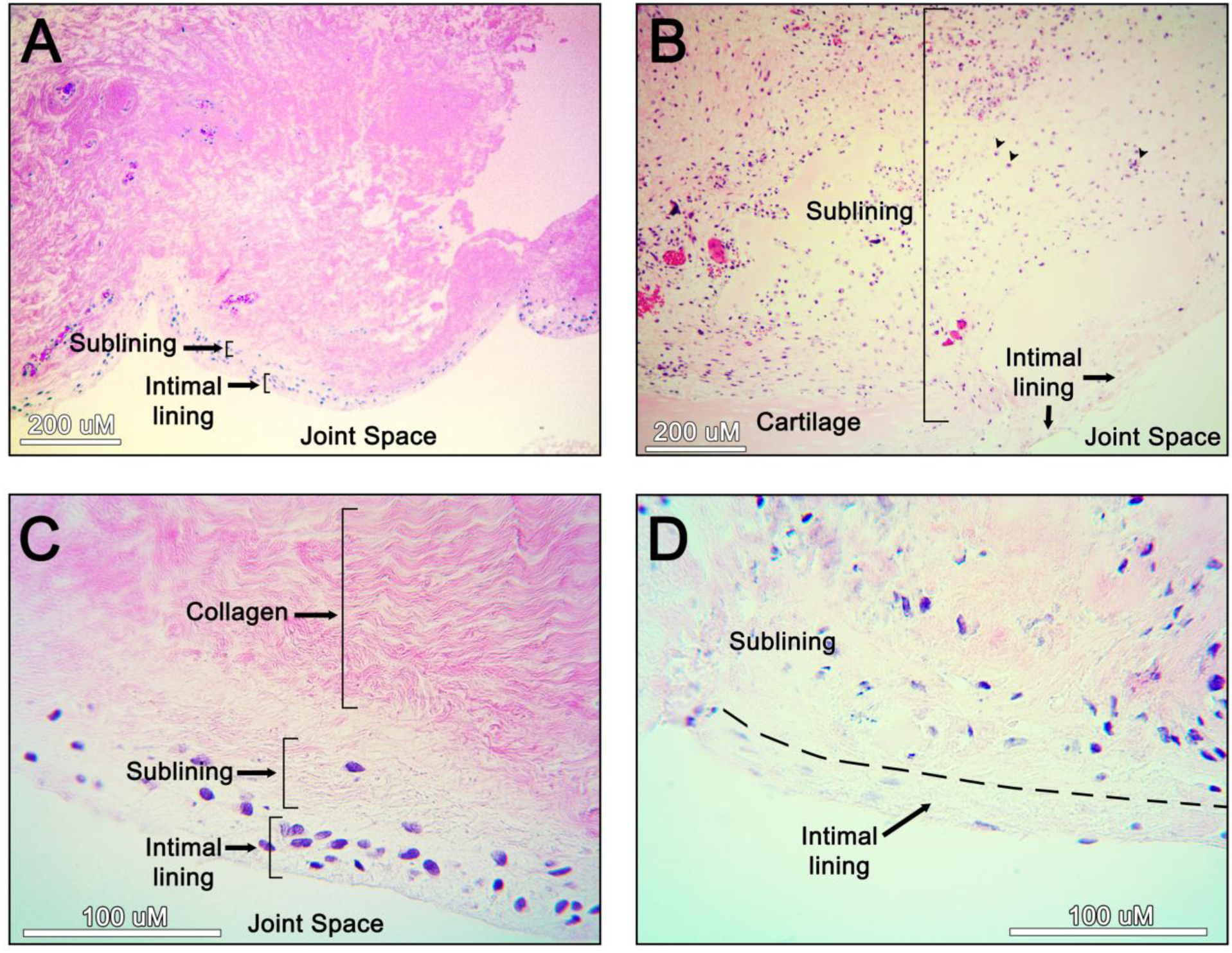
*In vivo* negative (A, C, same patient, 10X and 40X respectively) and positive (B, D, second patient, 10X and 40X respectively) controls. Negative control patient displays fibrosis, but positive control (chronic bacterial infection) patient has evident dysfunction of synovium with cartilage formation, evident migration of inflammatory cells into the sublining, and thinning/loss of cellularity of the initial lining (B, D).

**Supplementary Figure 3:**
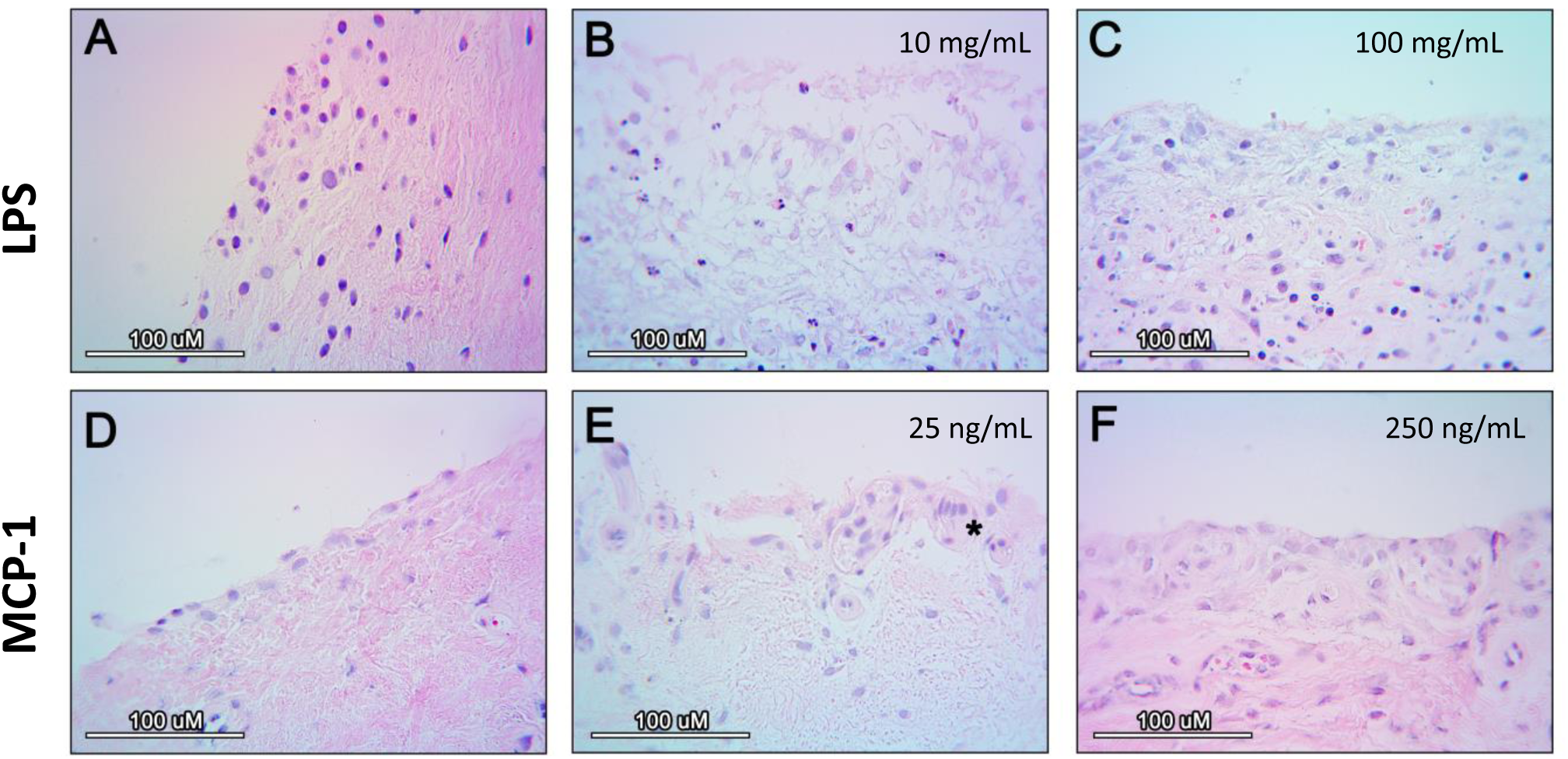
disruption of the synovial intimal lining and migration of inflammatory cells at 24 hours is dependent upon stimulus dose. A-C are from same patient, D-F are from a second patient, all at 24 hours.

**Supplemental Figure 4:**
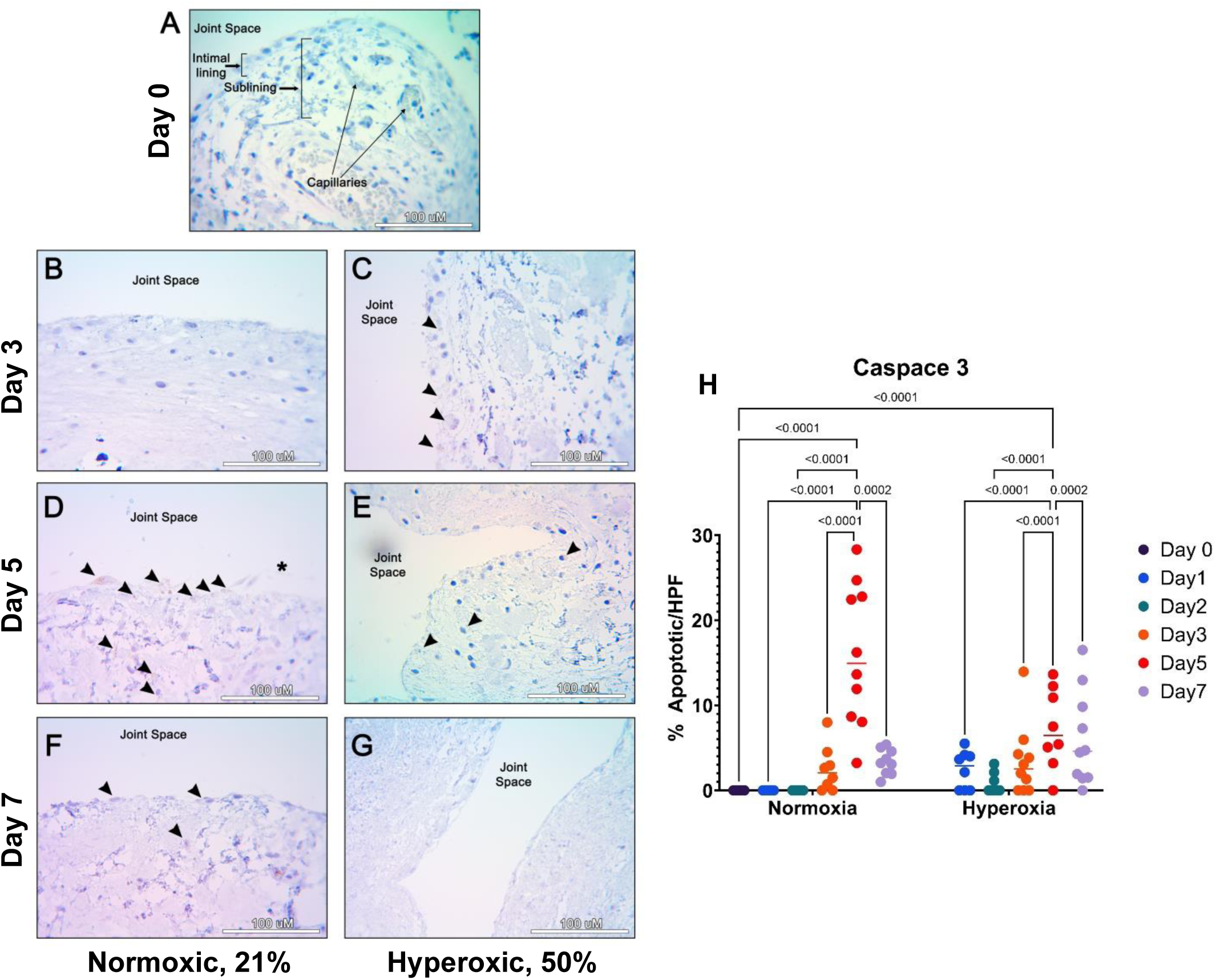
Caspace-3 IHC and quantification for explant culture out to 7 days in normoxic (B, D, F) and hyperoxic (C, E, G) conditions. Arrowheads point out positive IHC for Caspace-3. Apoptosis became evident in normoxic conditions on day 3 of culture, compared to day 1 for hyperoxic conditions. Two-way ANOVA with Tukey’s *post-hoc* correction (H).

**Supplementary Figure 5:**
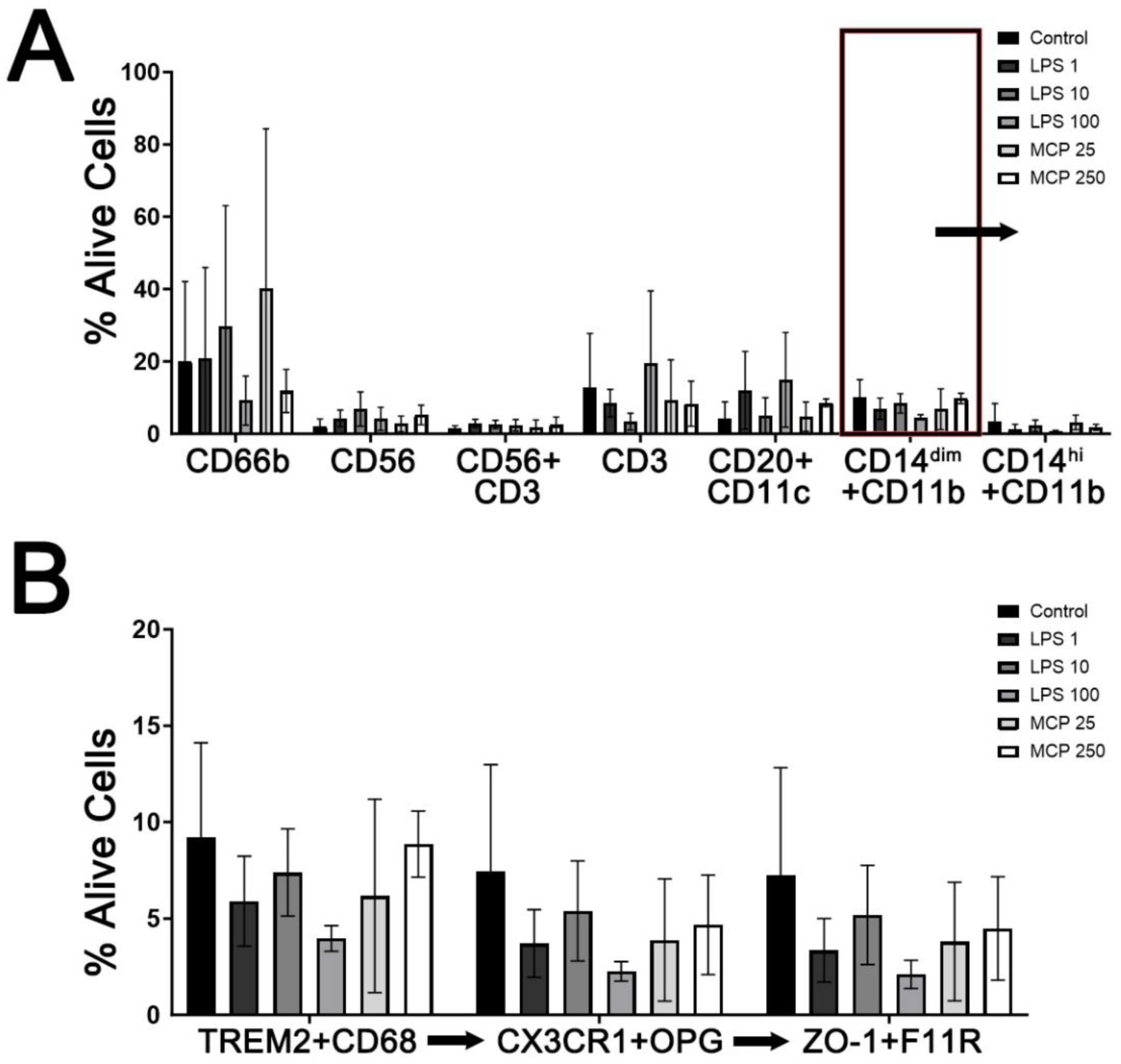
Evaluation of migratory cell populations in the bottom transwell. Neutrophils (CD66b), NK (CD56), NKT (CD3 and CD56), T cells (CD3), B cells or Dendritic Cells (CD20 and CD11c, same fluorophore), and CD14 high or dim myeloid (CD11b) cells (**A**). Sequential analysis of the CD14^dim^ myeloid population demonstrating nearly all CD14^dim^ macrophages are positive for OPG, CX3CR1, ZO-1, and F11R (**B**). LPS in µg/mL, MCP-1 in ng/mL.

**Supplemental Figure 6:**
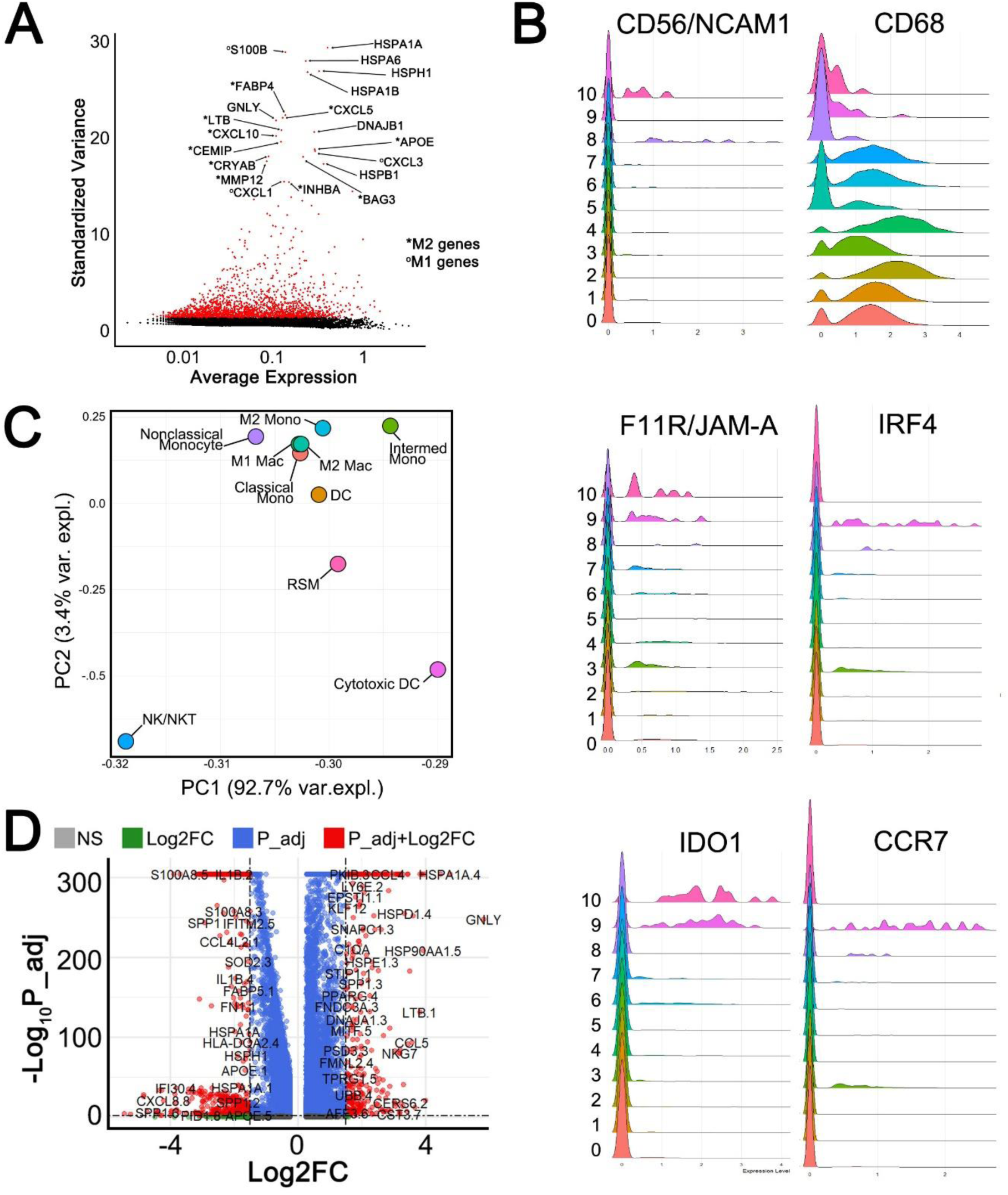
Evaluation of scRNA-seq data. Variance analysis with top 20 genes labeled (A). Ridge plots of NK, M2, and RSM associated genes (B). Principle component analysis of all 10 clusters not including dead cells (C). Volcano plot of most up- and down-regulated genes, where minimum Log2FC was 1.5 for significance given homogeneity of sample (D).

**Supplemental Figure 7:**
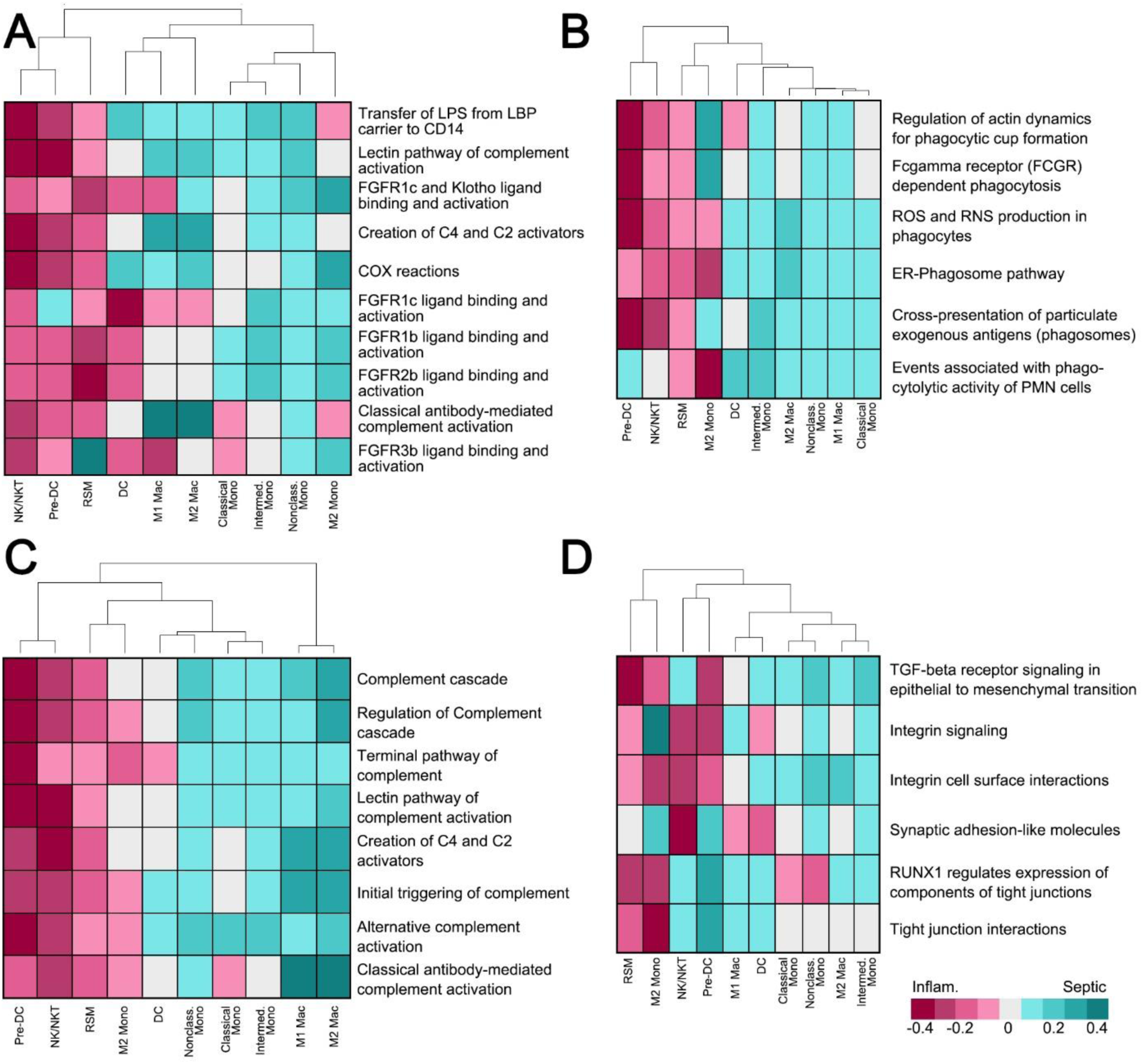
GSA for top 10 immune-relevant pathways (A), phagocytosis (B), complement (C), and adhesion-related (D).

**Supplemental Figure 8:**
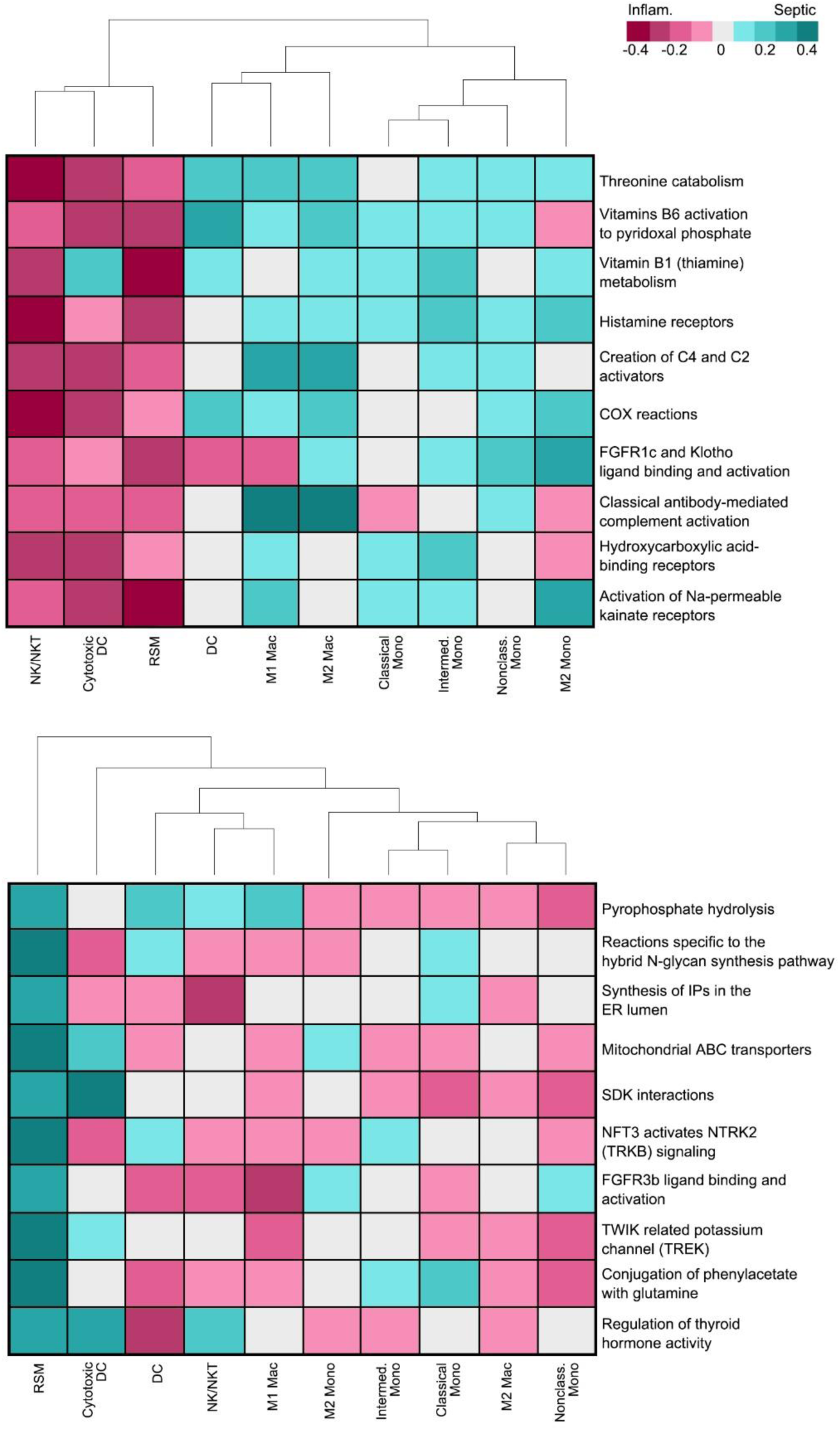
GSA for RSMs: top ten genes upregulated in inflammatory arthritis (top) and septic arthritis (bottom).

